# Kindlin-2 inhibits TNF/NF-κB-caspase 8 pathway in hepatocytes to maintain liver development and function

**DOI:** 10.1101/2022.07.13.499909

**Authors:** Huanqing Gao, Yiming Zhong, Liang Zhou, Sixiong Lin, Xiaoting Hou, Zhen Ding, Yan Li, Qing Yao, Huiling Cao, Xuenong Zou, Di Chen, Xiaochun Bai, Guozhi Xiao

## Abstract

Inflammatory liver diseases are a major cause of morbidity and mortality worldwide; however, underlying mechanisms are incompletely understood. Here we show that deleting the focal adhesion protein Kindlin-2 in hepatocytes using the *Alb-Cre* transgenic mice causes a severe inflammation, resulting in premature death. Kindlin-2 loss accelerates hepatocyte apoptosis with subsequent compensatory cell proliferation and accumulation of the collagenous extracellular matrix, leading to massive liver fibrosis and dysfunction. Mechanistically, Kindlin-2 loss abnormally activates the tumor necrosis factor (TNF) pathway. Blocking activation of the TNF signaling pathway by deleting TNF receptor or deletion of caspase 8 expression in hepatocytes essentially restores liver function and prevents premature death caused by Kindlin-2 loss.Finally, of translational significance, adeno-associated virus mediated overexpression of Kindlin-2 in hepatocytes attenuates the D-galactosamine and lipopolysaccharide-induced liver injury and death in mice. Collectively, we establish that Kindlin-2 acts as a novel intrinsic inhibitor of the TNF pathway to maintain liver homeostasis and may define a useful therapeutic target for liver diseases.

## Introduction

The liver is a multifunctional organ that plays critical roles in regulation of metabolism and detoxification and maintenance of the whole-body homeostasis[1]. Liver is comprised of parenchymal cells and non-parenchymal cells. The former includes hepatocytes and endothelial cells; the latter includes hepatic stellate cells (HSC) and Kupffer cells (KC). Abnormal activation of the TNF signaling stimulates inflammation and apoptosis, which are intertwined during the development and progression of liver diseases [2–5]. TNFα exerts its actions through binding and activating two different types of receptors, TNF receptor 1 (TNFR1) and TNFR2 [6]. It is known that NF-κB, a maor TNFR downstream effector, plays a pivotal role in promoting inflammatory liver diseases, such as viral and alcoholic hepatitis and fulminant hepatic failure [7]. Thus, it is critical to keep the TNF/NF-κB signaling activity under strict check to maintain normal organogenesis and function. However, key signal(s) that restrict abnormal TNF/TNFR activation in liver under physiological conditions and if and how alterations of these signals contribute to development and progression of inflammatory liver diseases are incompletely understood.

Kindlin proteins are key components of the focal adhesion (FA) assembly and contain the FERM domain, which is responsible for binding to and activating integrins to regulate the cell-extracellular matrix (ECM) adhesion, migration and signaling [8–13]. Mammals have three Kindlin proteins, i.e., Kindlin-1, Kindlin-2 and Kindlin-3, which are encoded by *Fermt1, Fermt2* and *Fermt3* genes, respectively. Kindlin-1 and Kindlin-3 are expressed primarily in epithelial and hematopoietic cells, respectively, while Kindlin-2 is widely expressed. Up-regulation of Kindlin-2 expression is implicated in a number of pathological processes, such as tumor formation, progression and metastasis [14, 15]. Global deletion of Kindlin-2 expression causes early embryonic lethality at E7.5 in mice [13]. Furthermore, Kindlin-2 and other FA related proteins are widely involved in the control of the development and function of skeleton, kidney, heart and other organs and tissues[16–32] through both integrin-dependent and integrin-independent mechanisms.

In this study, by utilizing biochemical and genetic mouse approaches, we establish that Kindlin-2 acts as a potent inhibitor of the TNF/NF-κB-caspase 8 pathway in hepatocytes and plays an important role in maintaining normal liver development and function.

## Results

### Kindlin-2 deletion in hepatocytes causes acute liver injury and premature death in mice

To investigate whether Kindlin-2 plays a role in liver development, we deleted its expression by crossing the floxed Kindlin-2 mice (*Kindlin-2*^*fl/fl*^) with *Alb-Cre* transgenic mice. The cross-breeding gave rise to genotypes at the expected mendelian ratio at birth. Results from quantitative real-time RT-PCR (qRT-PCR) and western blotting analyses revealed that the levels of Kindlin-2 mRNA and protein were decreased in livers, but not brain, heart, lung, kidney and spleen, in KO mice, compared to those in control littermates (Supplementary Figure 1). Shockingly, all KO mice (>30 mice) died at ages between 4 and 5 weeks (Figure 1a). At 4 weeks of age, both body and liver weights of KO mice were slightly, but significantly, decreased compared to those of control littermates (Figure 1b, c). At this age, livers from KO mice became coarsened and granular (Figure 1d, top panels). Massive ascites was observed in all KO mice but not in control mice (Figure 1d, bottom panels). KO mice displayed a splenomegaly (Figure 1e) and severe cholestasis (Figure 1f). Compared to sera from control mice, sera from KO mice were more yellowish in color (Figure 1g) and displayed elevated serum levels of both direct and indirect bilirubin (Figure 1h), suggesting that the animals suffered from cholestasis. KO mice displayed an osteopenic phenotype with significant reductions in both trabecular and cortical bone volume (Supplementary Figure 2).

**Figure 1.**
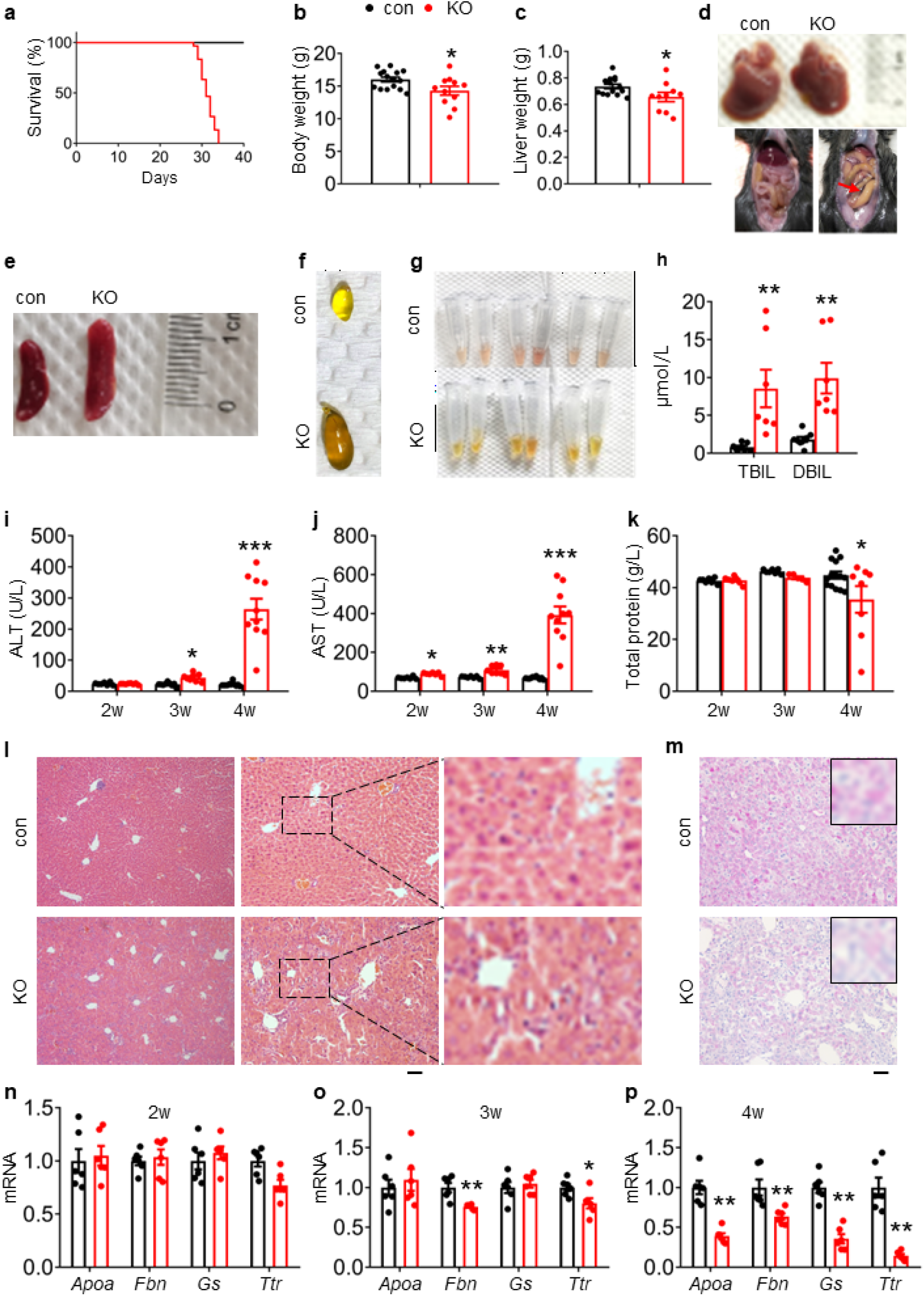
Kindlin-2 loss in hepatocytes causes liver injury and premature death in mice. (**a**) Survival curve of control and KO mice (*N* = 30 mice/group). (**b**) Body weight of 4-week-old control and KO mice (*N* = 15 for control mice, *N* =11 for KO mice). (**c**) Liver weight of 4-week-old control and KO mice (*N* = 14 for control mice, *N* = 10 mice for KO mice). (**d**) Livers and gross appearance of 4-week-old control and KO mice. Red arrow indicates massive ascites in KO. (**e**) Spleens from 4-week-old control and KO mice. (**f**) Gallbladder from 4-week-old control and KO mice. (**g**) Serum appearance from 4-week-old control and KO mice. (**h**) Serum total bilirubin (TBIL) and direct bilirubin (DBIL) levels. (*N* = 8 for control mice, *N* = 7 mice for KO mice). (**i, j**) Serum aminotransferase (ALT and AST) activity in 2-, 3- and 4-week-old control and KO mice. (**k**) Serum total protein levels in 2-, 3- and 4-week-old control and KO mice. (**l**) H/E staining of liver sections of 4-week-old control and KO mice Scale bars,100 μm. (**m**) Periodic acid-Schiff (PAS) staining for liver glycogen at 4 weeks of age. Scale bars,100 μm. (**n-p**) Quantitative real-time RT-PCR (q-RT-PCR) analysis of expression of liver genes in 2-, 3- and 4-week-old control and KO mice (*N* = 6 mice/group). The results are shown as means ± SEM. *\*P* < 0.05, \*\**P* < 0.01, vs control.

We further analyzed serum biochemistry in KO mice and age- and sex-matched littermates. The levels of both serum alanine transaminase (ALT) and aspartate transaminase (AST) activities were increased in KO mice relative to those in control mice, starting as early as 2 weeks of age with most dramatic increases at 4 weeks of age for both enzymes (Figure 1i, j). The levels of serum protein were reduced in a time-dependent manner (Figure 1k). The level of serum low density lipoprotein (LDL) was increased and that of serum high density lipoprotein (HDL) was decreased in KO relative to control mice (Supplementary Figure 3a, b). The levels of serum total cholesterol (TCH) and ammonia were increased in KO versus control mice (Supplementary Figure 3c, d). Results from hematoxylin and eosin (H/E) staining of liver sections of 4-week-old control and KO mice revealed that Kindlin-2 loss caused massive inflammatory cell infiltration in KO liver, especially in areas surrounding the hepatic sinusoidal vessels (Figure 1l). Kindlin-2 loss reduced the glycogen accumulation, as determined by Periodic acid-Schiff (PAS) staining (Figure 1m) and the mRNA expression levels of liver-related genes, including those encoding glutamine synthetase (Gs), apolipoprotein A-I (Apoa), fibrinogen b (Fbn) and transthyretin (Ttr) (Figure 1n-p).

Collectively, these results demonstrate that Kindlin-2 loss in hepatocytes causes a severe liver failure and damages in multiple, resulting in premature death.

### Kindlin-2 loss induces dramatic hepatocyte apoptosis and stimulates proliferation of both biliary cells and hepatic stellate cells in liver

A progressive increase in hepatocyte apoptosis was observed in KO mice, as demonstrated by TUNEL staining (Figure 2a, b and Supplementary Figure 4). Consistently, immunohistochemical (IHC) staining showed that the immune reactivity of caspase 3 protein was dramatically increased in KO relative to control livers (Figure 2c). Western blotting revealed that level of the pro-apoptotic Bax protein was time-dependently increased in KO livers compared to that in control livers (Figure 2d). In contrast, expression of the anti-apoptotic Bcl-2 protein was largely down-regulated in a time-dependent manner by Kindlin-2 deficiency (Figure 2d). Western blotting showed that the levels of the proliferation cell nuclear antigen (Pcna) and cyclin D1 proteins were increased in KO livers compared with those in control livers at 3 and 4 weeks of ages, but not at 2 weeks of age (Figure 2e). Results from IHC staining of control and KO liver sections using an anti-Ki67 antibody revealed an increase in cell proliferation in KO liver tissue (Figure 2f, g). Kindlin-2 loss apparently increased the number of cytokeratin 19 (CK19)-expressing cells, i.e., the biliary/progenitor cells (Figure 2h). Immunofluorescence (IF) staining revealed that the number of desmin-expressing cells, i.e., hepatic stellate cells (HSC), was dramatically increased in KO relative to that in control liver tissues (Figure 2i, j).

**Figure 2.**
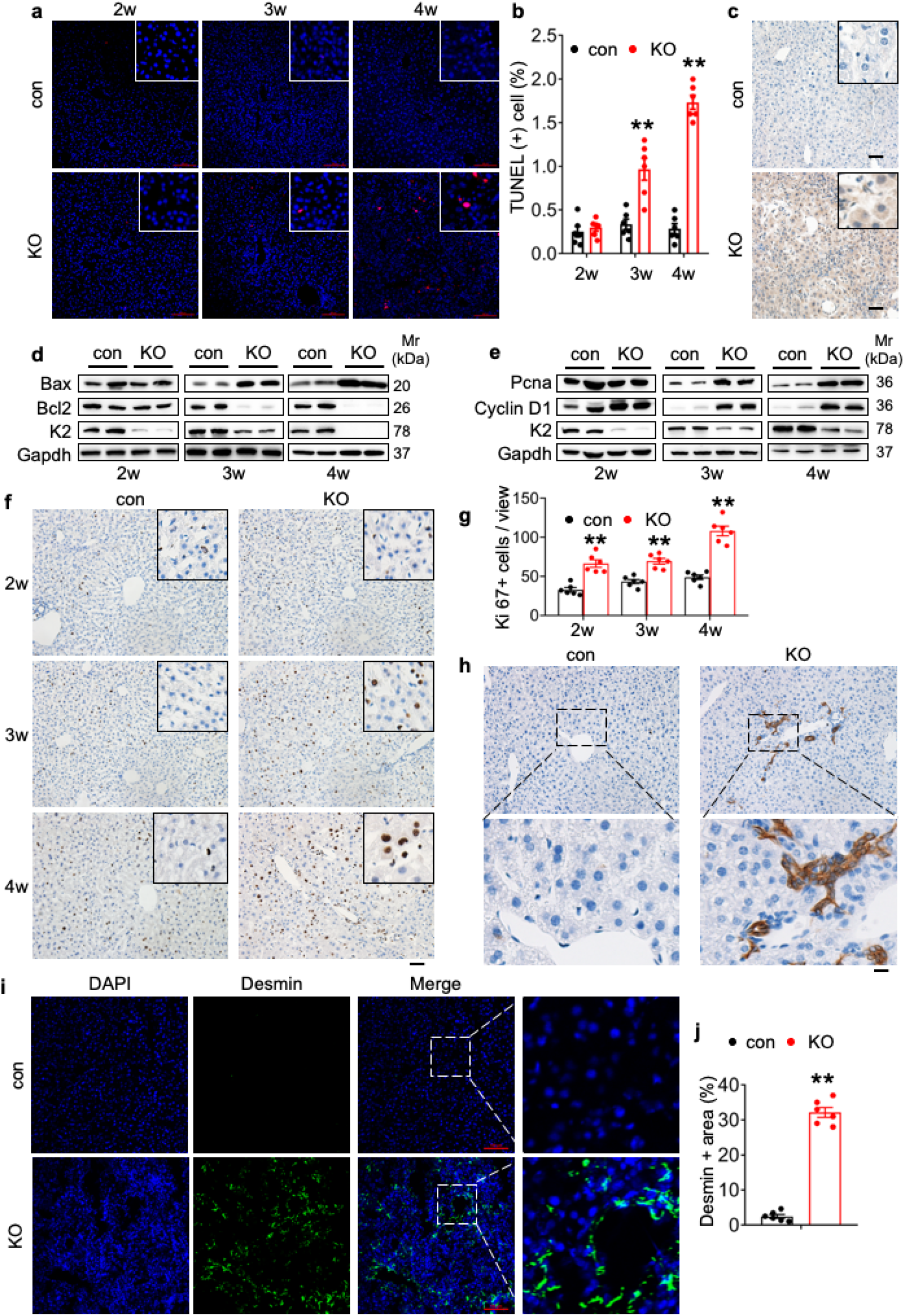
Kindlin-2 loss causes dramatic hepatocyte apoptosis followed by enhanced proliferation of both biliary cells and hepatic stellate cells. (**a**) TUNEL staining. Scale bars,100 μm. (**b**) Quantification of TUNEL-positive cells from 6 different fields from 6 mice per group. (**c**) Immunohistochemistry (IHC) staining for Caspase 3 was performed on paraffin liver sections of 4-week-old control and KO mice. Scale bars,100 μm. (**d**,**e**) Western blotting. Liver tissue lysates from 2-, 3- and 4-week-old control and KO mice were subjected to western blotting analysis for expression of the indicated proteins. Gapdh was used as a loading control. (**f**,**g**) Ki67 staining. Liver sections from of 2-, 3- and 4-week-old control and KO mice were subjected to Ki67 staining. Scale bars,100 μm. Quantification of (f). (**h**) IHC staining. Liver sections from 4-week-old control and KO mice were subjected to IHC staining for CK19. Scale bars,100 μm. Representative images showing a dramatic increase in the number CK19-positive cells in KO liver. (**i, j**) Immunofluorescence (IF) staining. Scale bars,100 μm. Liver sections from 4-week-old control and KO mice were subjected to IF staining for Desmin, a marker for HSC. Representative images showing high expression of Desmin in KO liver. Quantification of (i) from 6 mice per group. The results are shown as means ± SEM. *\*P* < 0.05, \*\**P* < 0.01, vs control.

### Kindlin-2 loss stimulates collagen deposition and massive fibrosis in liver

A dramatic increase in collagen deposition was observed in livers of 4-week-old KO mice, as demonstrated by the Masson trichromatic and Sirius red staining (Figure 3a-d). Consistently, results from western blotting revealed that level of α-smooth muscle actin (α-Sma), a marker for fibrosis, was drastically increased in KO livers in a time-dependent manner (Figure 3e). IHC staining was also reveal that α-Sma was upregulated in KO mice (Figure 3f). In addition, the mRNA levels of fibrogenic genes, including those encoding collagen α-1(I) chain (Col1a1), transforming growth factor β1 (Tgfβ1), TIMP metallopeptidase inhibitor 1 (Timp1), actin α2 (Acta2) and collagen type VI, α1 (Col6a1), were up-regulated in KO mice relative to those in control mice (Figure 3g).

**Figure 3.**
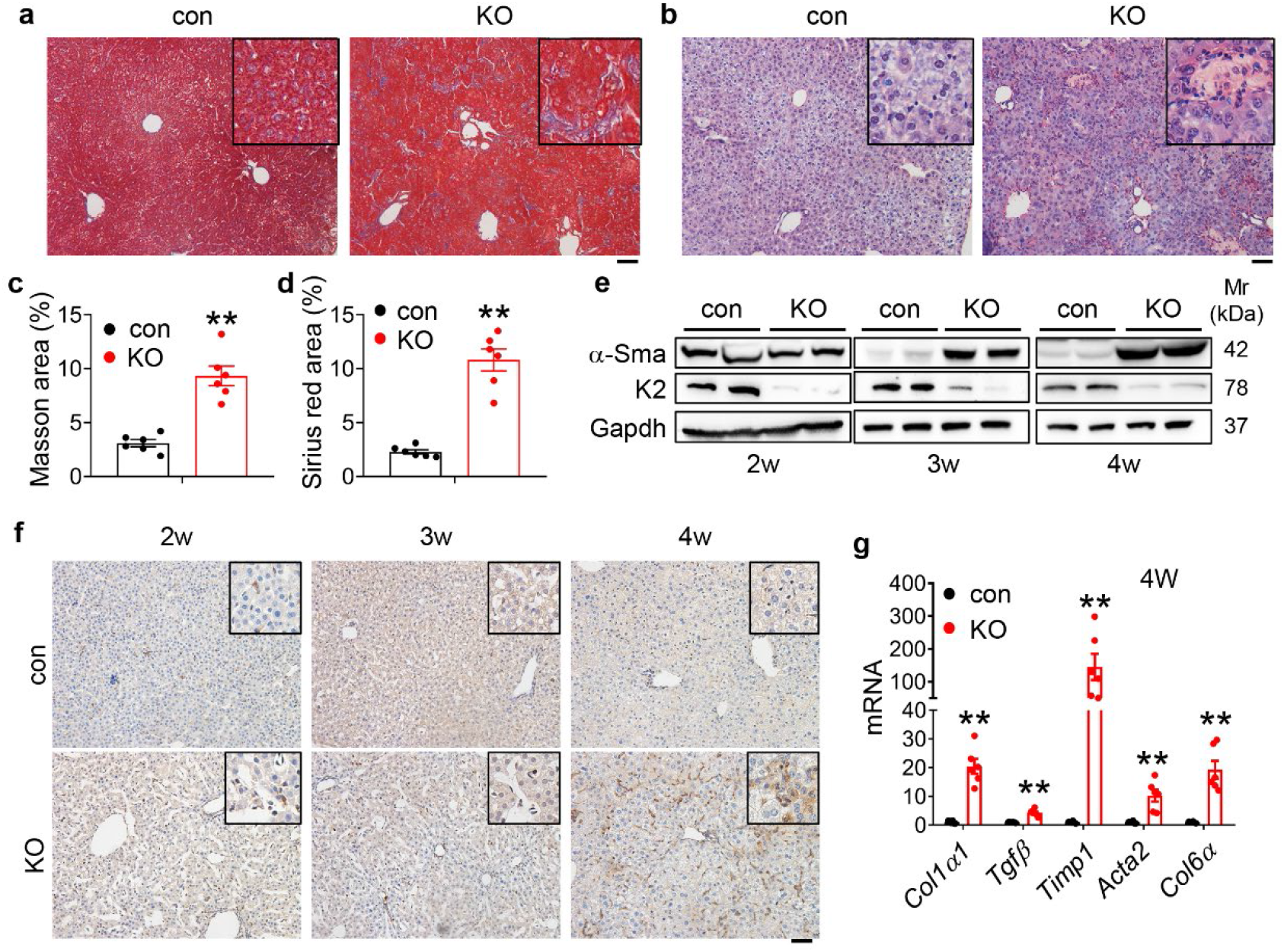
Kindlin-2 loss promotes collagen ECM deposition and fibrosis in liver. (**a**) Masson’s trichrome staining of liver sections of 4-week-old control and KO mice. Scale bars,100 μm. (**b**) Sirius red staining. Fibrillar collagen deposition in liver sections of 4-week-old control and KO mice was determined by Sirius red staining. Scale bars,100 μm. (**c, d**) Quantification of (a) and (b) from 6 mice per group. (**e**) Western blotting. Liver tissue extracts from 2-, 3- and 4-week-old control and KO mice were subjected to western blotting for expression of α-SMA and Kindlin-2. Gapdh was used as loading control. (**f**) IHC staining. Liver sections from 2-, 3- and 4-week-old control and KO mice were subjected to IHC staining using an antibody against α-SMA. Scale bars,100 μm. (**g**) qRT-PCR analysis. RNAs isolated from liver tissues of 4-week-old control and KO mice were subjected to qRT-PCR analysis for expression of fibrosis-related genes. (*N* = 6 mice/group). The results are shown as means ± SEM. \**P* < 0.05, \*\**P* < 0.01, vs control.

### Kindlin-2 loss greatly activates the TNF/NF-κB signaling pathway in vitro and in vivo

We further performed RNA-seq analyses from 4-week-old control and KO liver tissues and identified 6746 genes that exhibited statistically significant changes and differential expression (3496 up-regulated genes and 3250 down-regulated genes with log2FC>1.5). Kyoto Encyclopedia of Genes and Genome (KEGG) analysis revealed that Kindlin-2 loss impacted multiple important pathways. Kindlin-2 loss activated TNF signaling pathway, apoptosis and MAPK pathways in hepatocytes (Figure 4a, b). Consistent with result from the RNA-seq analysis, the serum level of tumor necrosis factor α (Tnfα) was dramatically elevated in 4-week-old KO mice compared to that in control littermates (Figure 4c). Furthermore, the mRNA levels of Tnfα was increased in a time-dependent manner in KO livers compared to those in control livers (Figure 4d). IHC staining of F4/80 revealed massive macrophage infiltration in KO but not control liver tissues (Figure 4e). The numbers of circulating neutrophils and other myeloid cells (e.g., monocytes) were increased in KO versus control mice (Figure 4f-h and Supplementary Figure 5).

**Figure 4.**
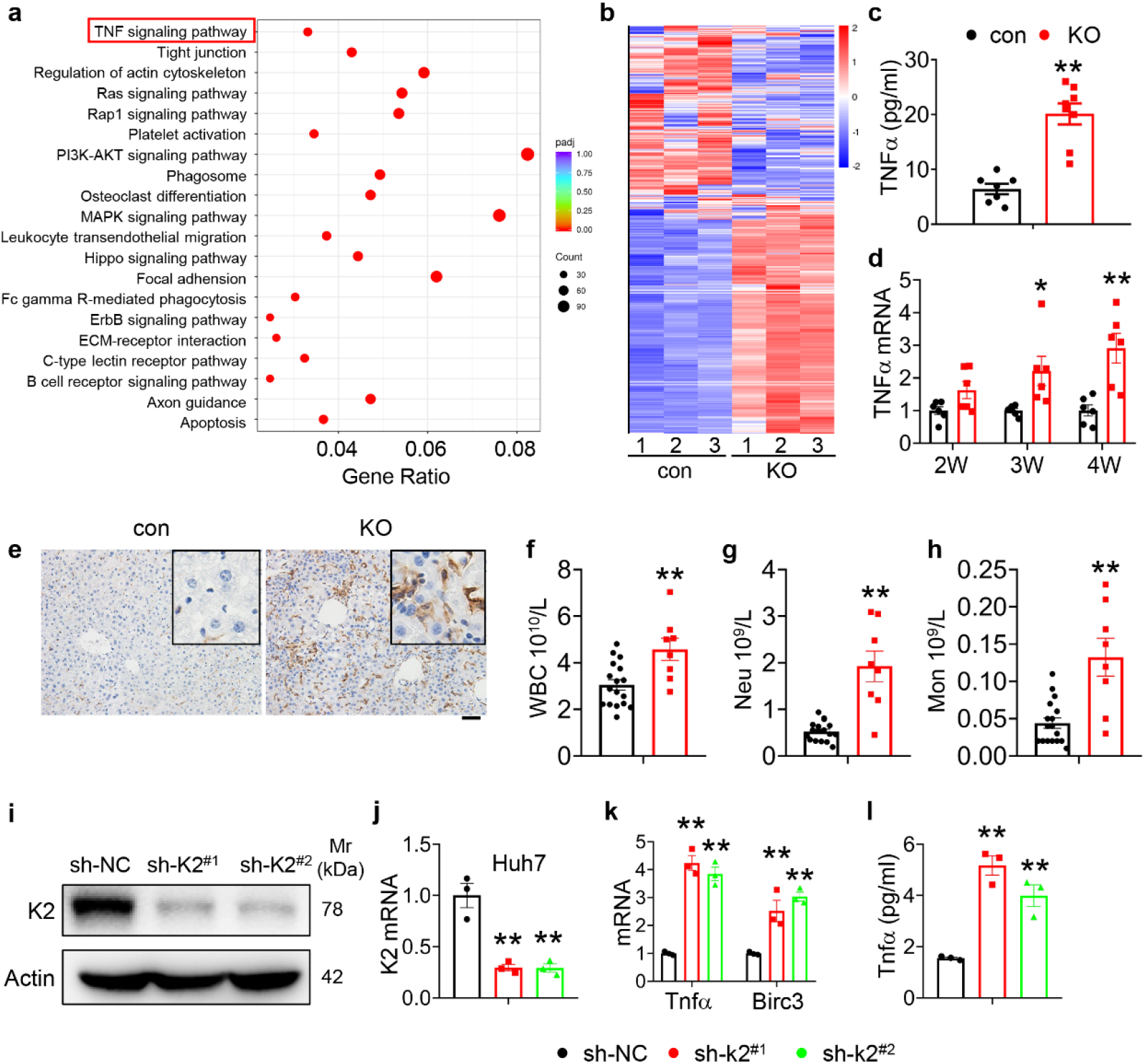
Kindlin-2 loss activates the TNF/NF-κB signaling pathway. (**a**) KEGG analysis showing the up-regulated pathways in the 4-week-old KO mice versus control mice. (**b**) Heatmap represents genes with >1.5-fold upregulation or >1.5-fold downregulation in K2-deficient liver compared with control liver. (**c**) Sera from 4-week-old control and KO mice were subjected to ELISA for Tnfα protein (*N* = 7-8 mice/group). (**d**) RNAs isolated from liver tissues of 2-, 3- and 4-week-old control and KO mice were subjected to qRT-PCR analyses for *Tnfα* genes (*N* = 6 mice/group). (**e**) 4-week-old control and KO mouse liver sections were subjected to IHC staining using an anti-F4/80 antibody to determine macrophage infiltration. Scale bars,100 μm. (**f-h**) Complete blood count were assessed on 4-week-old mice. WBC: white blood cell, Neu: neutrophil, Mon: monocyte. *N* = 17 for control mice and *N* = 8 for KO mice. (**i, j**) Kindlin-2 knockdown. qPCR analyses and western blotting were performed to detect Kindlin-2 expression in Huh7 cells treated with lentiviruses-expressed control shRNA (sh-NC) and two different Kindlin-2 shRNAs (sh-K2^(#1)^, sh-K2^(#2)^). (**k**) qPCR analysis of *Tnfa* and *BircI3* mRNAs in sh-NC- and sh-K2-treated HepG2 cells. (**l**) ELISA. The levels of Tnfa protein in media of sh-NC- and sh-K2-treated Huh7 cultures were assayed using an ELISA kit. Each sample was tested at least in triplicate independent cell preparations.The results are shown as means ± SEM. \**P* < 0.05, \*\**P* < 0.01, vs control.

To define the function of Kindlin-2 in hepatocyte in vitro, we knocked down its expression in Huh7 and HepG2 cells. The levels of Kindlin-2 mRNA and protein were dramatically reduced by Kindlin-2 shRNA (sh-K2) in both Huh7 (Figure 4i, j) and HepG2 cells (Supplementary Figure 6a, b). The mRNA levels of Tnfα and Birc3 (Baculovial IAP repeat containing 3) were up-regulated by Kindlin-2 knockdown in Huh7 (Figure 4k) and HepG2 cells (Supplementary Figure 6c). ELISA showed a dramatic increase in TNF protein in supernatants of the sh-K2 Huh7 (Figure 4l) and HepG2 cultures (Supplementary Figure 6d). Collectively, these results illustrate that ablation of Kindlin-2 in hepatocyte resulted in upregulation of TNF signalling pathway.

### TNFR ablation reverses liver lesions and lethality caused by Kindlin-2 deficiency

To determine whether abnormal activation of the TNF signaling plays a key role in mediating the liver damage and death caused by Kindlin-2 deficiency, we globally deleted the expression of the tumor necrosis factor receptors in Kindlin-2 KO mice and determined its effects. We used a breeding strategy by crossing the *Tnfr^−/−^* mice, in which both *Tnfrsf1a* and *Tnfrsf1b* are globally deleted, with the *Alb-Cre; Kindlin-2*^*fl/+*^ mice to generate the Tnfr-deficient Kindlin-2 KO mice (referred to as KO/*Tnfr^−/−^* hereafter) (Supplementary Figure 7). *Tnfr^−/−^* mice were purchased from Shanghai Biomodel Organism Science & Technology Development Co., Ltd. The KO/*Tnfr^−/−^* mice were normal at birth with the expected Mendelian ratio. Strikingly, deletion of *Tnfr* genes completely prevented the premature death of the Kindlin-2 KO mice. In contrast to the fact that all Kindlin-2 KO mice died between 4-5 weeks of ages, no KO/*Tnfr^−/−^* mice (>30 mice) died during this period of time and beyond (Figure 5a). The levels of serum ALT and AST were significantly lower in KO/*Tnfr^−/−^* mice than in Kindlin-2 KO mice (Figure 5b, c). As shown in (Figure 5d), deletion of *Tnfr* genes rescued the liver damage in Kindlin-2 KO mice. The increases in collagen deposition, macrophage infiltration, cell proliferation, fibrosis and apoptosis caused by Kindlin-2 deficiency were markedly reversed by *Tnfr* genes deletion (Figure 5e-n).

**Figure 5.**
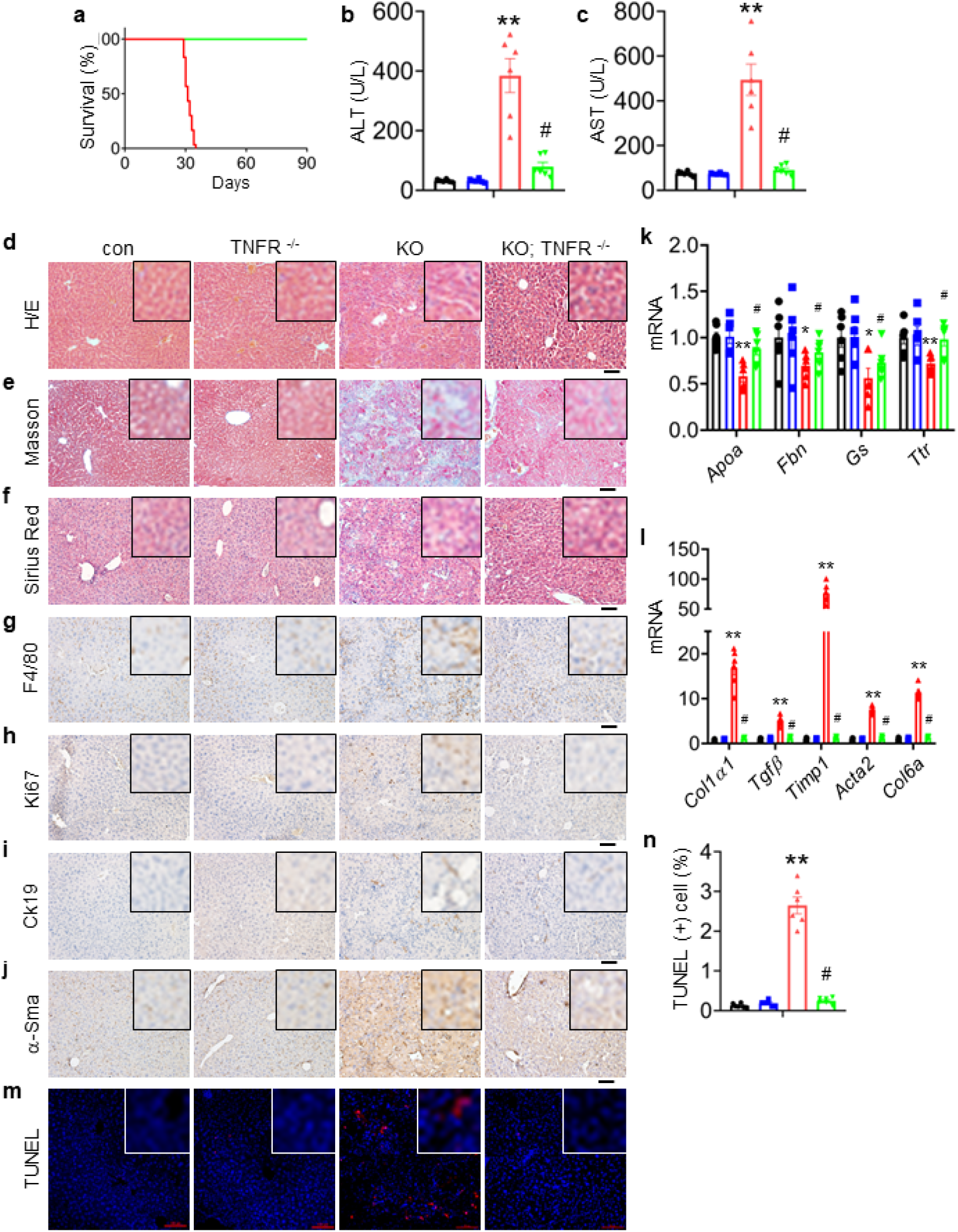
Global *Tnfr* genes deletion rescues the liver injury and lethality induced by Kindlin-2 loss. (**a**) Survival curve of KO and KO/*Tnfr^−/−^* mice. *N* = 30 mice in KO group, *N* = 33 for KO/*Tnfr^−/−^* group. (**b, c**) Serum ALT and AST levels in control, KO, *Tnfr^−/−^* and KO/*Tnfr^−/−^* mice. *N* = 6 mice/group. (**d-f**) Liver histology. Liver sections from mice of the indicated genotypes were subjected to H/E staining (d), Masson’s trichrome staining (e) and Sirius Red staining (f). Scale bars,100 μm. (**g-j**) IHC staining of liver sections from mice of the indicated genotypes for expression of F4/80 (g), Ki67 (h), Ck19 (i) and α-Sma (j). Scale bars,100 μm. (**k, l**) qRT-PCR analysis. RNAs isolated from liver tissues of the indicated genotypes were subjected to qRT-PCR analysis for expression of the indicated genes. *N* = 6 mice/group. (**m, n**) TUNEL staining of the indicated genotypes and quantification. *N* = 6 mice/group. Scale bars,100 μm. The results are shown as means ± SEM. \**P* < 0.05, \*\**P* < 0.01, vs control. ^#^*P* < 0.05, vs KO.

### Caspase 8 deletion in hepatocytes attenuates liver lesions and prevents lethality of Kindlin-2 KO mice

It is known that abnormal inflammation causes cell death and that caspase 8 is a major initiator in the death receptor-mediated apoptosis. We therefore next determined whether Kindlin-2 loss causes hepatocyte death and thereby liver lesions and lethality by activating the caspase 8 dependent apoptotic pathway. To this end, we generated mice lacking both Kindlin-2 and caspase 8 in hepatocytes (referred to as KO/*Cas8^−/−^* hereafter). Shockingly, deletion of caspase 8 in hepatocytes essentially prevented premature death of Kindlin-2 KO mice (Figure 6a). The levels of serum ALT and AST were significantly lower in KO/*Cas8^−/−^* mice than in Kindlin-2 KO mice (Figure 6b, c). Deletion of caspase 8 restored the liver damage in Kindlin-2 KO mice (Figure 6d). The increases in macrophage infiltration, cell proliferation, fibrosis and apoptosis caused by Kindlin-2 deficiency were markedly attenuated by caspase 8 deletion (Figure 6e-l).

**Figure 6.**
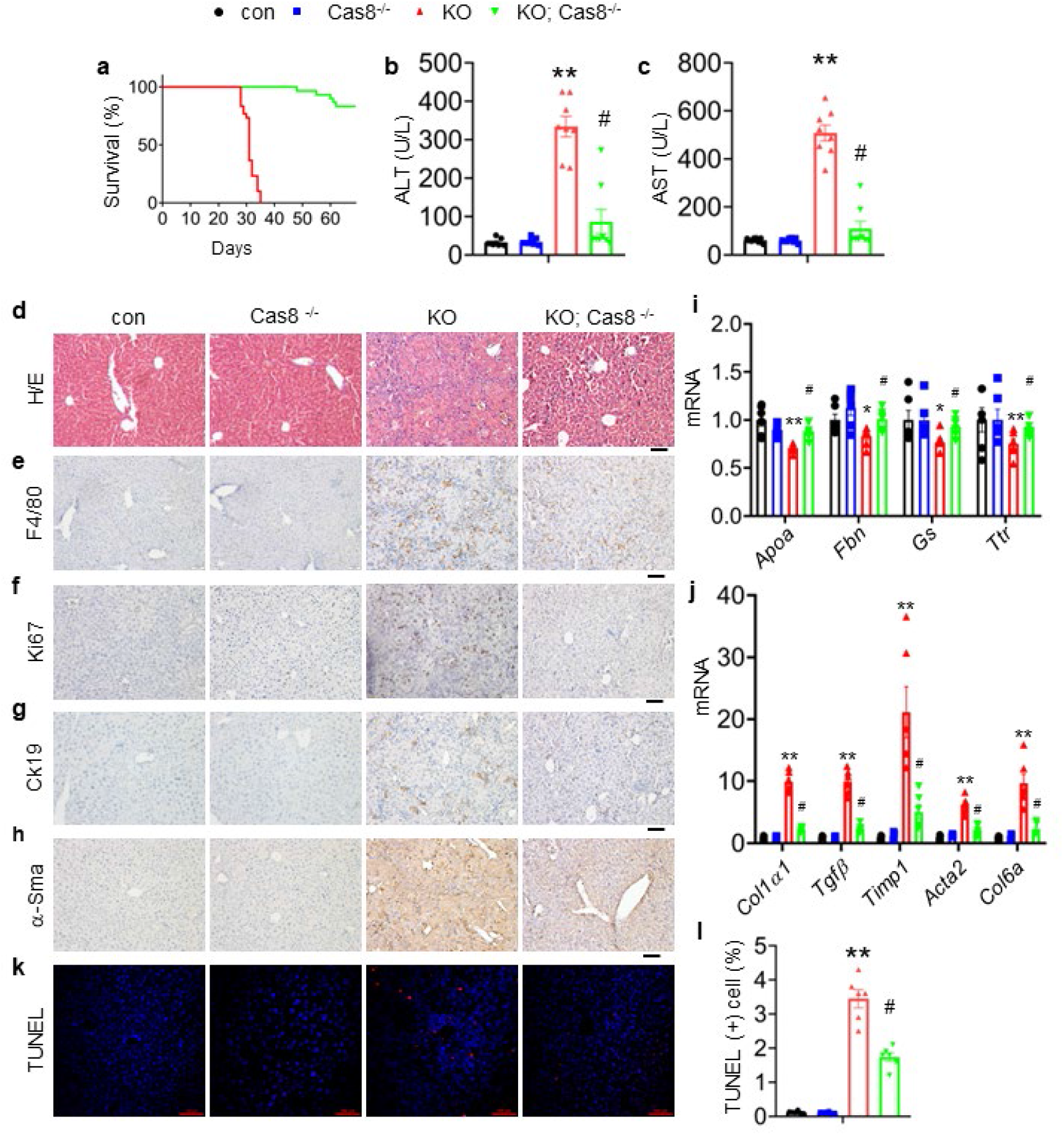
Caspase 8 deletion in hepatocytes rescues the liver lesions and lethality caused by Kindlin-2 deficiency. (**a**) Survival curve of KO and KO/*Cas8^−/−^* mice. *N* = 30 mice/group. (**b, c**) Serum ALT and AST levels in control, KO, *Cas8^−/−^* and KO/*Cas8^−/−^* mice. *N* = 8 mice/group. (**d**) Liver histology. Liver sections from mice of the indicated genotypes were subjected to H/E staining, Scale bars,100 μm. (**e-h**) IHC staining of liver sections from mice of the indicated genotypes for expression of F4/80 (e), Ki67 (f), Ck19 (g) and α-Sma (h). Scale bars,100 μm. (**i, j**) qRT-PCR analysis. RNAs isolated from liver tissues of the indicated genotypes were subjected to qRT-PCR analysis for expression of the indicated genes. *N* = 6 mice/group. (**k, l**) TUNEL staining of the indicated genotypes and quantification. *N* = 6 mice/group. Scale bars,100 μm. The results are shown as means ± SEM. \**P* < 0.05, \*\**P* < 0.01, vs control. ^#^*P* < 0.05, vs KO.

### AAV8-mediated overexpression of Kindlin-2 in hepatocytes attenuates the GalN/LPS-induced liver injury and death

We finally investigated the effect of Kindlin-2 on the liver injury induced by D-galactosamine (GalN)/lipopolysaccharide (LPS). Immunoblotting analysis detected a dramatic decrease in Kindlin-2 accumulation in the liver tissues of C57BL/6 after administration with GalN/LPS (Figure 7a, b). Next, 8-week-old C57BL/6 mice were first injected via tail vein with adeno-associated virus 8 (AAV8) (2×10^11^ particles/mouse) expressing Kindlin-2 (AAV8-K2) or GFP (AAV8-GFP). After 21d, mice were treated with GaIN/LPS for 5h or a longer period of time (for survival curve experiment) (Figure 7c). The level of Kindlin-2 protein was markedly increased in livers of mice injected with AAV8-K2 (Figure 7d, e). As expected, GalN/LPS resulted in acute death of all mice around 14-15 hours of the treatment. Kindlin-2 overexpression (OE) significantly extended the life span of mice compared with control mice injected with AAV8-GFP (Figure 7f). Kindlin-2 OE improved the gross liver appearance in GalN/LPS-treated mice (Figure 7g). Kindlin-2 OE significantly decreased the levels of serum ALT and AST in GalN/LPS-treated mice (Figure 7h, i). Results from H/E staining revealed that histological damages in liver caused by GalN/LPS were markedly ameliorated by Kindlin-2 OE (Figure 7j). Furthermore, Kindlin-2 OE decreased GalN/LPS-induced hepatocyte apoptosis, as measured by immunohistochemical staining of Caspase-3 and TUNEL staining of liver sections (Figure 7k-m).

**Figure 7.**
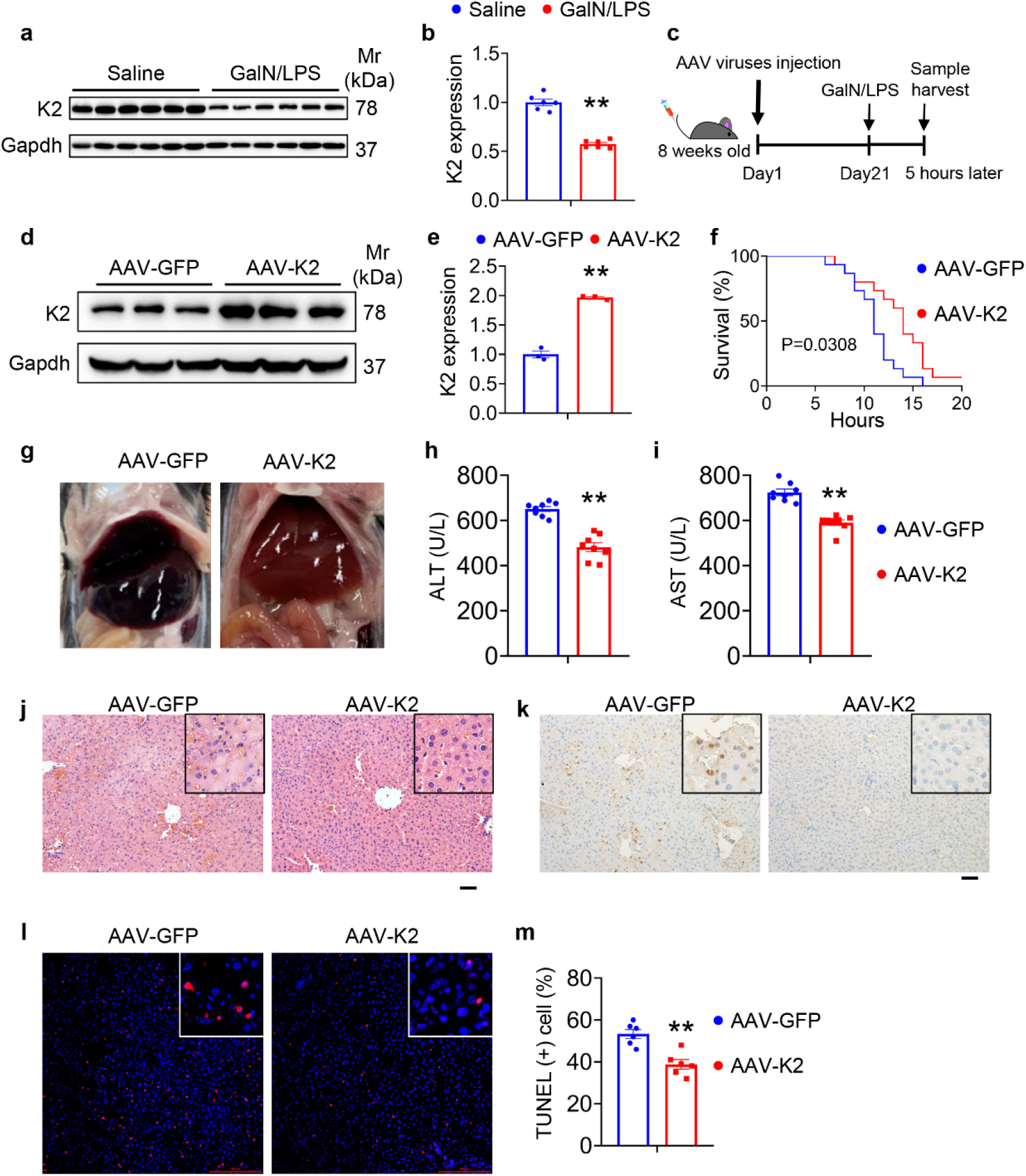
Overexpression of Kindlin-2 ameliorates D-GalN/LPS-induced liver injury and death. (**a, b**) Eight-week-old C57BL/6 mice were intraperitoneally injected with D-GaIN/LPS (GalN, 700 mg/kg body weight, LPS, 3 mg/kg body weight) or PBS as controls for 5 h. Liver tissues were collected and subjected to western blotting for expression of Kindlin-2. Quantitative data (a). *N* = 6 mice/group. (**c**) Experimental design. (**d, e**) Western blotting. Eight-week-old C57BL/6 mice were first injected via tail vein with adeno-associated virus 8 (AAV8) (2×10^11^ particles/mouse) expressing Kindlin-2 (AAV8-K2) or GFP (AAV8-GFP). After 21d, mice were then treated with D-GaIN/LPS for 5 h. Protein extracts were then prepared from livers and subjected to western blotting for expression of Kindlin-2. Quantitative data (d). Gapdh was used as a loading control. *N* = 3 mice per group. (**f**) Survival curve. Mice were treated as in (c), followed by observation for death. *N* = 15 mice per group. (**g**) Gross liver appearance. Mice were treated as in (c). (**h, i**) ELISA assays. Serum ALT (f) and AST (g). Mice were treated as in (c). *N* = 8 mice per group. (**j**) H/E staining in liver sections. Mice were treated as in (c). (**k**) IHC staining of caspase-3 in liver sections. Mice were treated as in (c). (**l, m**) TUNEL staining of liver sections. Quantitation data (m). Mice were treated as in (c). *N* = 6 mice per group. \*\**P* < 0.01, vs AAV-GFP.

## Discussion

In this study, we demonstrate a novelt role of the key focal adhesion protein Kindlin-2 in regulation of liver development and function. Kindlin-2 loss in hepatocytes causes an acute liver failure and premature death in mice. We demonstrate that Kindlin-2 loss dramatically activates the TNF/NF-κB signaling pathway, massive inflammation and hepatocyte death with subsequent stimulation of proliferation of both biliary cells and hepatic stellate cells, leading to ECM accumulation and fibrosis in liver. We further demonstrate that liver damages and lethality caused by Kindlin-2 deficiency are largely attenuated or completely rescued by either global ablation of TNFR or hepatocyte-selective deletion of caspase 8 expression. We finally demonstrate that adeno-associated virus mediated overexpression of Kindlin-2 in hepatocytes attenuates the D-galactosamine/LPS-induced liver injury and death in mice.

In this study, we demonstrate that Kindlin-2 expressin in hepatocytes is essential for liver development and function. Kindlin-2 deletion in hepatocytes using the Alb-Cre transgenic mice causes acute liver failure and lethality of 100% penetration in mice at ages between 4 and 5 weeks. Our results suggest that abnormal activation of the TNF/NF-κB signaling pathway plays a major role in mediation of the liver lesions and lethality caused by Kindlin-2 deficiency. This notion is supported by the following lines of in vitro and in vivo evidences. First, results from our RNA-seq analyses from 4-week-old control and KO liver tissues reveal that TNF signaling pathway is activated in KO liver. Second, the levels of *Tnfα* mRNA in liver and serum level of TNFα protein were dramatically elevated in 4-week-old KO versus control mice. Consistently, massive macrophage infiltration is observed in KO but not control liver tissues, suggesting increased inflammation. Third, in vitro studies show that siRNA knockdown of Kindlin-2 expression increases the mRNA levels of *Tnfα* and *Birc3* in Huh7 and HepG2 cells. Fourth, the increases in serum ALT and AST, hepatocyte apoptosis, ECM deposition, inflammatory infiltration, proliferation of both biliary cells and hepatic stellate cells, and liver fibrosis caused by Kindlin-2 deficiency were largely reversed by *Tnfr* genes deletion. Most importantly, *Tnfr* genes deletion completely rescues the lethality caused by Kindlin-2 deficiency. These findings also indicate the necessity to keep the TNF/NF-κB signaling pathway in hepatocytes under precise control in order to maintain normal liver development and function as well as the whole-body metabolic homeostasis. We demonstrate that Kindlin-2 plays a pivotal role in this regard.

Kindlin-2 loss induces massive hepatocyte apoptosis. The expression of the pro-apoptotic Bax protein is up-regulated and that of the anti-apoptotic Bcl2 protein is down-regulated in mutant liver. A number of apoptotic hepatocytes are observed throughout the mutant liver. Strikingly, genetic ablation of caspase 8 expression in hepatocytes essentially rescues the liver lesions and lethality caused by Kindlin-2 loss. These results suggest that the caspase 8 dependent death receptor-mediated apoptotic pathway plays a critical role in mediating the liver damages and lethality caused by Kindlin-2 loss. Abnormal inflammation induced by Kindlin-2 loss should largely contribute to hepatocyte apoptosis in the mutant mice.

Notably, Kindlin-2 loss greatly promotes proliferation of both biliary cells and hepatic stellate cells and stimulates accumulation and deposition of excessive collagenous ECM, which leads to liver fibrosis in the mutant mice.

Of translational significance, AAV8-mediated expression of Kindlin-2 in liver significantly blocks the D-GalN/LPS-induced liver inflammation and death in mice. In the furture, it would be important to investigate whether Kindlin-2 loss plays a role in the pathogenesis of human inflammatory liver diseases.

In summary, we establish that the focal adhesion protein Kindlin-2 is a potent intrinsic inhibitor of the TNF/NF-κB-caspase 8 inflammatory pathway and plays an important role in the maintenance of normal liver development and function. We may define a novel therapeutic target for inflammatory liver diseases.

## Materials and Methods

### Animal study

Generation of *Kindlin-2*^*fl/fl*^ mice was previously described[27]. To delete Kindlin-2 expression in hepatocyte, we first bred the *Kindlin-2*^*fl/fl*^ mice with the *Alb-Cre* transgenic mice[33], which were kindly provided by Dr. Yan Li of Southern University of Science and Technology, and obtained the *Kindlin-2*^*fl/+*^; *Alb-Cre* mice. Further intercrossing of the *Kindlin-2*^*fl/+*^; *Alb-Cre* mice with *Kindlin-2*^*fl/fl*^ mice generated the *Kindlin-2*^*fl/fl*^; *Alb-Cre* mice, the hepatocyte conditional Kindlin-2 knockout mice (referred to as KO). The Cre-negative floxed Kindlin-2 mice (*Kindlin-2*^*fl/fl*^) were used as controls in this study. *Tnfrsf1a^−/−^* and *Tnfrsf1b^−/−^* mice was obtained from the Shanghai Model Organism. *Caspase 8*^*fl/fl*^ mice were used as indicated[34]. All mice used in this study have been crossed with normal C57BL/6 mice for more than 10 generations. Mice were maintained in 12-hour light/dark cycles, with unrestricted access to food and water. All animal experiments were approved and conducted in the specific pathogen free (SPF) Experimental Animal Center of Southern University of Science and Technology (Approval number: 20200074).

### Biochemical measurements

Blood and tissues were collected from mice under anesthesia with isoflurane. Blood collected was allowed to clot for 2 h at room temperature and then centrifuged to collect sera. Serum total cholesterol (TCH), aspartate aminotransferase (AST), alanine aminotransferase (ALT) and albumin were measured with commercial kits (Shensuoyoufu, Shanghai, China).

### Histological analyses

Tissues were fixed in 4% PFA and then embedded in paraffin. Serial 5-μm paraffin sections were used for H/E staining using our standard protocols. Masson trichrome staining and Sirius Red staining were performed as described previously[35]. Immunohistochemistry was performed using paraffin sections according to a protocol previously described[19]. In brief, samples were deparaffinized and rehydrated, followed by antigen retrieval in 10 mM sodium citrate. Blocking and staining were performed in antibody diluent with background-reducing components (DAKO). Samples were incubated with primary antibodies as listed in supplementary table, followed by appropriate secondary antibodies. Images were obtained using a light microscope equipped with a digital camera.

### Immunofluorescence staining

Immunofluorescence (IF) staining was performed as previously described[19]. Briefly, liver frozen sections (10 μm thickness) were prepared using a Leica cryostat, fixed in 4% paraformaldehyde for 30 min, blocked for 2 h with 5% normal goat serum supplemented with 1% BSA, and incubated with the indicated antibodies at 4°C overnight. The sections were incubated with appropriate fluorescence-labelled secondary antibodies and analyzed by immunofluorescence microscopy.

### Cell culture in vitro

HepG2 (cat# SCSP-510) and Huh7 (cat# TCHu182) cells were purchased from the Cell Bank of the Chinese Academy of Sciences and cultured in DMEM supplemented with 10% fetal bovine serum, 1% penicillin, and streptomycin in a 5% CO2 incubator at 37 °C.

### Western blot analysis

For western blotting, total protein samples were extracted from tissues or cells and 30 μg of protein samples were separated on a 10% sodium dodecyl sulfate polyacrylamide gel electrophoresis gel and transferred to a polyvinylidene fluoride (PVDF) membrane. Protein expression was visualized by incubating primary antibodies overnight at 4°C, followed by the corresponding secondary antibodies and developed using the enhanced chemiluminescence system (Bio-Rad, #1705040)[34, 36]. Antibodies information are described in Supplementary Table 1.

### RNA-seq data and GO/KEGG enrichment analyses

Total RNAs were extracted from livers of 4-week-old control and KO mice. The sequencing library was determined by an Agilent 2100 Bioanalyzer using the Agilent DNA 1000 Chip Kit (Agilent, #5067-1504). Differentially expressed genes (DEGs) were selected with *P* value < 0.05, log2FC > 1 or < −1, and FPKM > 5 in at least one condition. Volcano plot showing dysregulated genes between control and KO livers was generated using R programming. GO and KEGG enrichment analyses were performed on the DEGs using TBtools.

### Quantitative real-time RT-PCR analysis

Total RNA from tissues and cells was extracted with Trizol reagent (Invitrogen, #10296010) as described[37]. After total RNA isolation and cDNA synthesis, PCR amplification was performed with the SYBR Green PCR Master Mix (Bio-Rad, #1725200). The mRNA expression levels of the target genes were normalized to expression of the glyceraldehyde 3-phosphate dehydrogenase (*Gapdh*). Each sample was tested at least in triplicate and repeated using three independent cell preparations. Primer sequences are listed in Supplementary Table 2 and Supplementary Table 3.

### TUNEL assay

Liver frozen sections were fixed with 4% paraformaldehyde and subjected to TUNEL staining using an In Situ Cell Death Detection Kit (Beyotime Biotechnology, #C1089), following manufacturer’s procedure.

### Enzyme-linked immunosorbent assay (ELISA)

After collection of sera from mice and supernatants from cultured hepatocytes, TNFα (R&D System, #MTA00B and DTA00D) and Il1β (R&D System, #MLB00C) levels were measured by ELISA kit according to manufacturer’s instruction.

### Statistical analysis

The sample size in mouse experiments of this study was determined based on our previous experience. Mice were randomly grouped in experiments in this study. IF, IHC and histology were performed and analyzed in a double-blinded way. The two-tailed unpaired Student’s *t* test (two groups) and one-way ANOVA (multiple groups), followed by Tukey’s post-hoc test, were used for statistical analyses using Prism GraphPad. The results are presented as means ± SEM (standard error of mean). *P* < 0.05 was considered statistically significant.

## Acknowledgements

The authors acknowledge the assistance of Core Research Facilities of SUS Tech. This work was supported by the National Key Research and Development Program of China Grants (2019YFA0906001), the National Natural Science Foundation of China Grants (81991513 and 81870532), Shenzhen Municipal Science and Technology Innovation Council Grants (JCYJ20180302174246105) and the Guangdong Provincial Science and Technology Innovation Council Grant (2017B030301018).

## Competing Interests

The authors declare no competing interests.

## Author Contributions

Study design: GX and HG. Study conduct and data collection and analysis: HG, YZ, LZ, XH, ZD, SL, YL, QY, HC, XZ, DC and XB. Data interpretation: GX and HG. Drafting the manuscript: GX and HG. GX and HG take the responsibility for the integrity of the data analysis.

## Data availability

All data generated or analysed during this study are included in the manuscript and supporting file; Source Data has been provided for Figures and Figure supplements.

## Notes

### Competing Interest Statement

The authors have declared no competing interest.

