## supplemental figures and tables for "Kindlin-2 inhibits TNF/NF-κB-caspase 8 pathway in hepatocytes to maintain liver development and function"

**Kindlin-2 inhibits the TNF/NF-κB pathway to maintain liver homeostasis**

Gao et.al

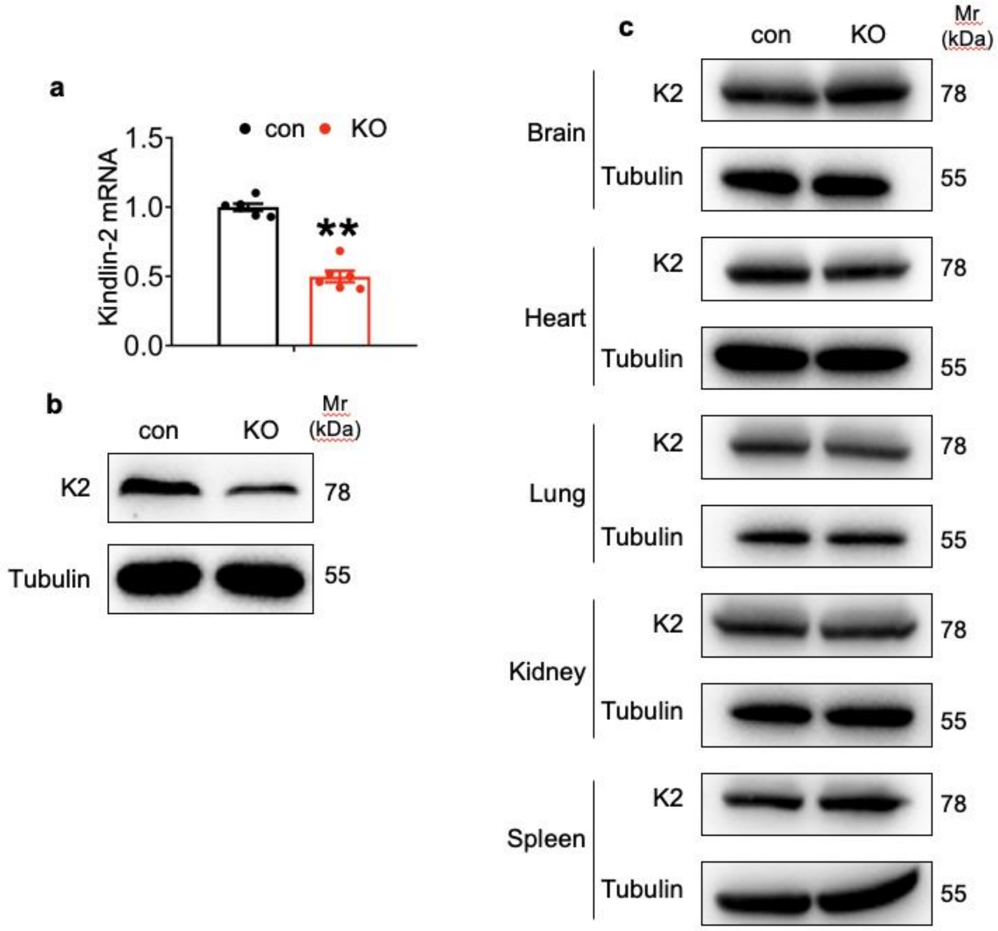


**Fig. S1. Deletion of Kindlin-2 in hepatocyte resulted in systemic dysfunction.** (**a**) Quantitative real-time RT-PCR (qRT-PCR) analysis. RNAs isolated from liver tissues of 4-week-old control and KO mice were subjected to qRT-PCR analysis (*N* = 6 mice/group). (**b**) Western blotting. Liver extracts from 4-week-old control and KO mice were subjected to western blotting analysis for expression of Kindlin-2. Tubulin was used as a loading control. (**c**) Western blotting. Protein expression of Kindlin-2 was determined by western blotting in the indicated tissues of 4-week-old control and KO mice. P**<0.01 vs control.

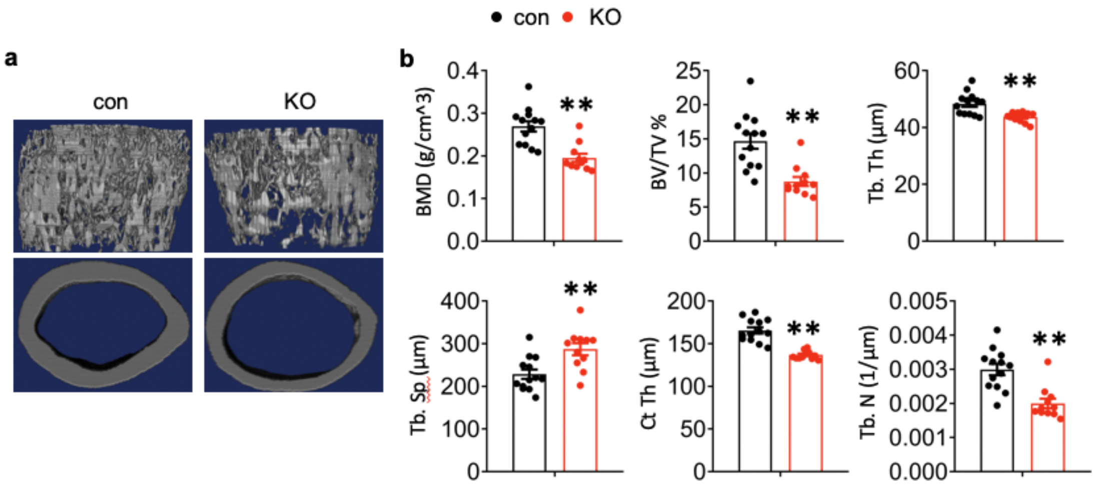


**Fig. S2. Deletion of Kindlin-2 in hepatocyte resulted in osteoporosis.** (**a**) Three-dimensional (3D) reconstruction from micro-computerized tomography (μCT) scans of femurs from con and KO mice. (**b**) Bone histomorphometric analyses (*N* =13 for control mice, *N* =11 for KO mice). P**<0.01 vs control.

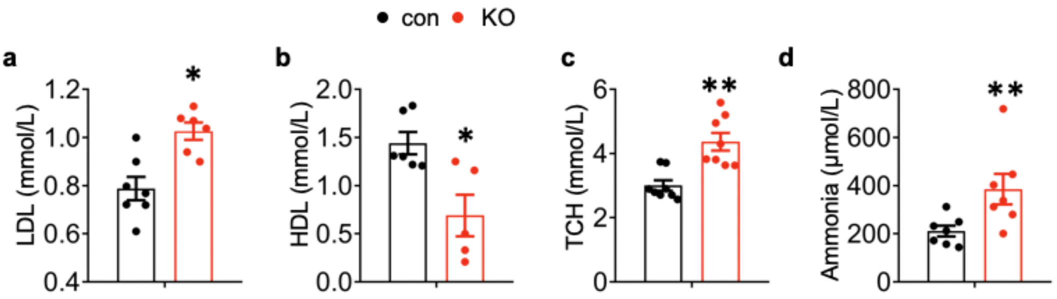


**Fig. S3 Metabolic parameters were assessed on 4-week-old mice (*N*= 7 mice per group).** (**a**) LDL: Low Density Lipoprotein, (*N* =7 for control mice, *N* =6 for KO mice). (**b**) HDL: High Density Lipoprotein. (*N* =6 for control mice, *N* =5 for KO mice). (**c**) TCH: total cholesterol. (*N* = 8 mice/group). (**d**) Ammonia level. (*N* = 7 mice/group). P*<0.05, P**<0.01 vs control.

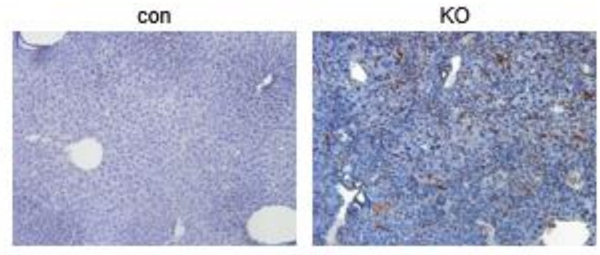


**Fig. S4. Loss of Kindlin-2 resulted in increased apoptosis.** TUNEL staining show increased number of apoptotic (yellow stain) cells in 4 weeks old KO liver. (magnification 200x).

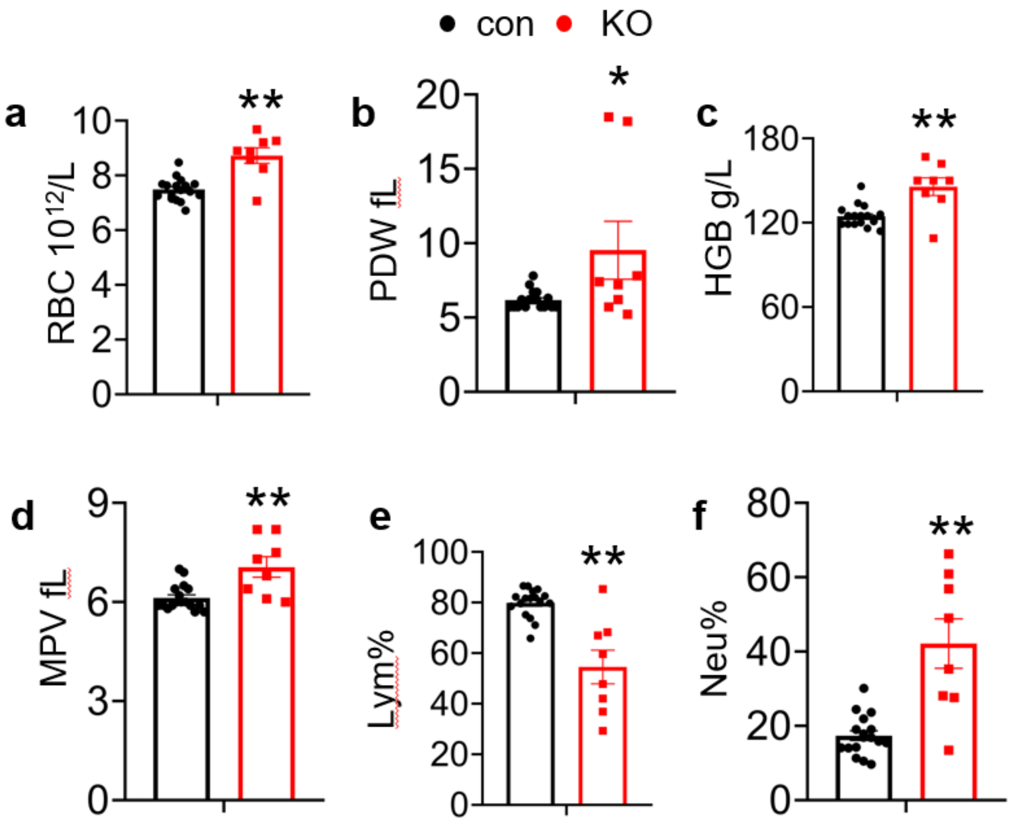


**Fig. S5 Complete blood count were assessed on 4-week-old mice (*N*= 17 for control mice and *N* = 8 for KO mice).** (**a**) RBC: red blood cell, (**b**) PDW: platelet distribution width, (**c**) HGB: hemoglobin, (**d**) MPV: mean platelet volume, (**e**) Lym: lymphocyte, (**f**) Neu: neutrophil, P*<0.05, P**<0.01 vs control.

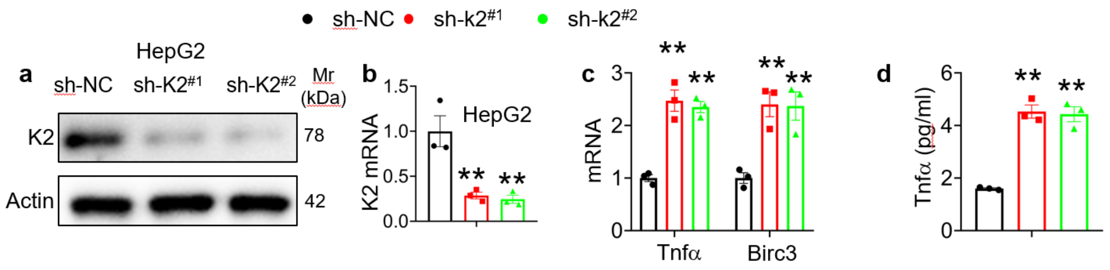


**Figure S6.** (a, b) Kindlin-2 knockdown. qPCR analyses and western blotting were performed to detect Kindlin-2 expression in HepG2 cells treated with lentiviruses-expressed control shRNA (sh-NC) and two different Kindlin-2 shRNAs (sh-K2$^{(\#1)}$, sh-K2$^{(\#2)}$). (c) qPCR analysis of *Tnfa* and *Bircl3* mRNAs in sh-NC- and sh-K2-treated HepG2 cells. (d) ELISA. The levels of Tnfa protein in media of sh-NC- and sh-K2-treated HepG2 cultures were assayed using an ELISA kit. P*<0.05, P**<0.01 vs shNC.

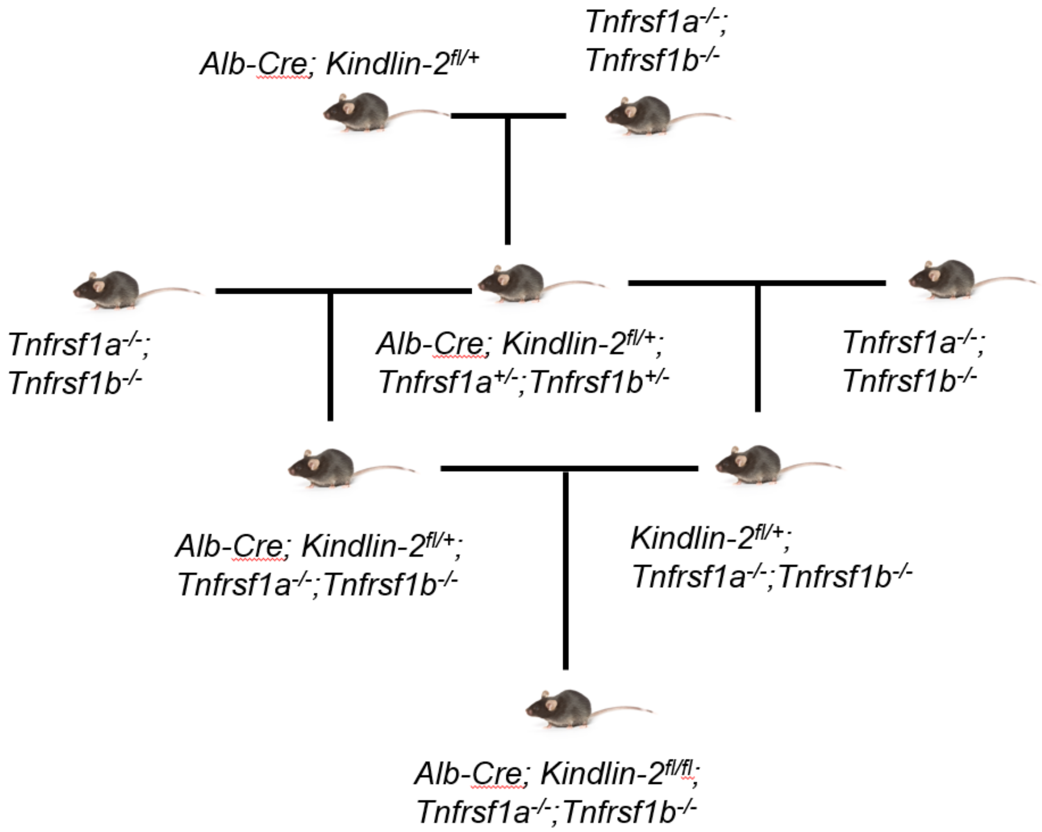


**Fig. S7** Breeding strategy to generate the *Alb-Cre; Kindlin-2*$^{fl/+}$*; Tnfrsf*$^{-/-}$ mice.

**Supplementary table1: antibody information**

| Name | Supplier | Cat no. | Application |
|---|---|---|---|
| Kindlin2 | Merck Millipore | MAB2617 | WB (1:1000) |
| Gapdh | ZSGB-BIO | TA-08 | WB (1:5000) |
| Bax | Cell Signaling technology | 5023S | WB (1:1000) |
| Bcl2 | Abcam | ab59348 | WB (1:1000) |
| Pcna | Abcam | Ab18197 | WB (1:1000) |
| Lamin B | EASYBIO | BE3191 | WB (1:1000) |
| Tubulin | Cell Signaling Technology | 2128 | WB (1:5000) |
| Cyclin D1 | Abcam | Ab16663 | WB (1:1000) |
| α-Sma | Absin | Abs120451 | WB (1:1000)<br>IHC (1:200) |
| Caspase3 | Sigma | MAB10753 | IHC (1:200) |
| F4/80 | Cell Signaling technology | 70076 | IHC (1:200) |
| Desmin | Cell Signaling technology | 5332 | IF (1:200) |
| Ck19 | Cell Signaling technology | 12434 | IHC (1:200) |
| Actin | ZSGB-BIO | TA-09 | WB (1:5000) |
| Ki67 | Cell Signaling technology | 12202 | IHC (1:200) |

**Supplementary table2: primer information for mouse**

| Gene | Forward | Reverse |
|---|---|---|
| Apoa | GGCACGTATGGCAGCAAGAT | TAGTCTCTGCCGCTGTCTTTGA |
| Ttr | CCATGAATTCGCGGATGTG | AGCCGTGGTGCTGTAGGAGTA |
| Fbn | GGATGGCAGCGTCGACTTT | TGGCAAGCCACAGTACTTCTTC |
| Tnfα | CCACGTCGTAGCAAACCACC | GATAGCAAATCGGCTGACGG |
| Col1a1 | TAGGCCATTGTGTATGCAGC | ACATGTTCAGCTTTGTGGACC |
| Col6a3 | ACTGGAACCACGGAAGTTCA | GTCACTTCCAACATCGAGGC |
| Tgfβ1 | GTGGAAATCAACGGGATCAG | ACTTCCAACCCAGGTCCTTC |
| Timp1 | AGGTGGTCTCGTTGATTTCT | GTAAGGCCTGTAGCTGTGCC |
| Acta2 | GTTCAGTGGTGCCTCTGTCA | ACTGGGACGACATGGAAAAG |
| Cs | TGGTCTGAAGTGCATTGAGGAG | CACCGGCAGAAAAGTCGTTG |
| Kindlin2 | TGGACGGGATAAGGATGCCA | TGACATCGAGTTTTTCCACCAAC |
| Gapdh | TTTCTTCTTGCCTTGGGAGA | AGTTCCGCACTTCATTCAGG |

**Supplementary table3: primer information for human**

| | | |
|---|---|---|
| BIRC3 | CTGGGCAGCAGGTTTACAA | GCATTCTTTGGATAGTAAAACACCA |
| KINDLIN-2 | GACCATGGCGGACAGTTCTT | TTCTCGCTGTTATCTGCTTGT |
| GAPDH | TCGGAGTCAACGGATTTGGT | TTCCCGTTCTCAGCCTTGAC |
| TNFα | TAGCCCATGTTGTAGCAAACC | GCTCTTGATGGCAGAGAGGA |
